# Selection for cell yield does not reduce overflow metabolism in *E. coli*

**DOI:** 10.1101/2021.05.24.445453

**Authors:** Iraes Rabbers, Willi Gottstein, Adam Feist, Bas Teusink, Frank J. Bruggeman, Herwig Bachmann

**Affiliations:** Systems Biology Lab, Vrije Universiteit Amsterdam, 1081 HV Amsterdam, The Netherlands; Department of Bioengineering, University of California, San Diego, La Jolla, CA, USA; NIZO Food Research, Ede, The Netherlands

**Keywords:** overflow metabolism, r/k selection, yield, emulsion culturing, metabolic strategy, cell size, experimental evolution

## Abstract

Overflow metabolism is ubiquitous in nature, and it is often considered inefficient because it leads to a relatively low biomass yield per consumed carbon. This metabolic strategy has been described as advantageous because it supports high growth rates during nutrient competition.

Here we experimentally evolved bacteria without nutrient competition by repeatedly growing and mixing millions of parallel batch cultures of *E. coli*. Each culture originated from a water-in-oil emulsion droplet seeded with a single cell. Unexpectedly we found that overflow metabolism (acetate production) did not change. Instead the numerical cell yield during the consumption of the accumulated acetate increased as a consequence of a reduction in cell size. Our experiments and a mathematical model show that fast growth and overflow metabolism followed by the consumption of the overflow metabolite, leads to a higher numerical cell yield and therefore a higher fitness compared to full respiration of the substrate. This provides an evolutionary scenario where overflow metabolism can be favourable even in the absence of nutrient competition.

## Introduction

When microbes compete, fast-growing strategies typically become dominant. While numerous microorganisms have the ability to respire during fast growth on high-quality carbon sources like glucose, many bacteria, yeasts and mammalian cells are known to not make full use of this capability, even when sufficient oxygen is available^1–4^. Instead of efficiently catabolizing the available carbon source to CO_2_, they produce metabolic by-products like acetate, lactate or ethanol at the expense of metabolic efficiency, i.e. biomass yield (gram biomass/mole carbon source). This phenomenon is termed overflow metabolism (Figure 1A) and has been suggested to be evolutionarily favourable at nutrient excess, because it supports the highest growth rates. It allows microorganisms that employ this strategy to outcompete fully respiratory cells^5–7^. When subjected to low substrate concentrations, cells grow at a lower rate and display full respiratory behaviour, e.g. in chemostats, which is associated with an increase in metabolic efficiency^1,2,6,8^.

**Figure 1.**
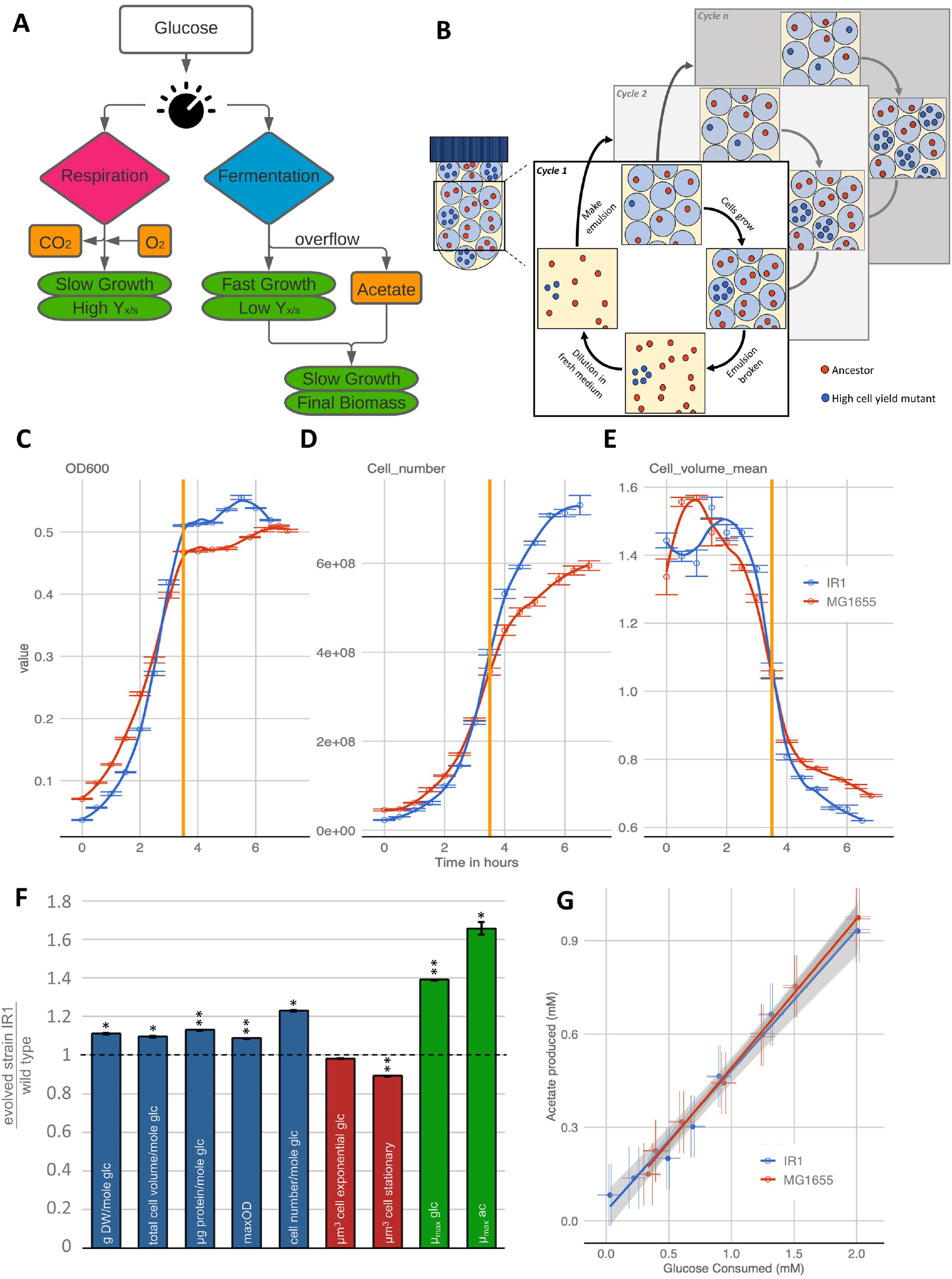
Cell propagation in emulsion droplets resulted in a mutant strain with increased cell yield, but not in a change of acetate production. (A) High biomass yield (Y_x/s_) is commonly associated with respiration. Upon depletion of the primary carbon source, growth on overflow metabolites leads to additional biomass formation. (B) Serial propagation of cells grown in emulsion droplets selects for mutants with an increased number of viable offspring per mole glucose (numerical cell yield). (C-E) After 53 transfers in emulsion a selected colony was compared to the wild type. In batch culture fast growth on glucose (depletion is indicated with the vertical orange line) is followed by slow growth on the formed acetate. During growth on acetate the cell number (counts/ml) increases much more than the optical density. Upon glucose depletion cell volume (μm3) decreases. The effect of cell number/volume is bigger for the evolved strain IR1 compared to the wild type MG1655. (F) A comparison of evolved strain IR1 normalized to the wildtype revealed significant increases in different biomass and cell yield measures (blue), as well as growth rate (green), while the cell size (red) decreased significantly during the second growth phase on acetate. Error bars represent standard deviations, one star indicates two-sided t-test p-value <0.05, two stars t-test p-value <0.005, n=3. For more details on the underlying data, see Supplementary Table S1 and Supplementary Figure S2. (G) The moles of acetate produced per moles of consumed glucose did not change significantly between the wild type and the selected strain IR1 (ANOVA; F=0.58, F-crit=4.26; p-value=0.45, α=0.05, n=3).

In nature many microbial species are in spatially structured environments where resources can be localized and the microbes undergo cycles of feast and famine. Their metabolic strategies have adapted to those conditions. Characteristic of such conditions are periods of nutrient excess followed by starvation, similar to batch cultures that are continued until complete depletion of the carbon sources. The winning strategy generates the highest number of viable offspring after a single cycle of feast and famine. Thus, microorganisms are faced with an “optimization problem”: the maximal number of viable offspring has to be produced in the shortest possible time given the available resources. In this context, metabolic inefficiency of overflow metabolism is generally considered to be the price for being fast.

As (growth) rate selection is associated with the occurrence of overflow metabolism, one would assume that yield selection should result in an increased metabolic efficiency, typically via enhanced respiratory metabolism. One way to ensure that growth rate competition between microorganisms is prevented, and selection of yield becomes possible, is by culturing individual cells in droplets of water-in-oil emulsions (Figure 1B). In this regime each cell gets its own “privatized” amount of substrate within a medium droplet, and does not compete for it with other genotypes, ruling out rate selection. The cells in a droplet can be allowed to grow until depletion of all carbon sources. When *L. lactis* was subjected to this protocol, the mutant cells that fixed had shifted their metabolism from inefficient homolactic to the more efficient mixed-acid fermentation. These mutants had a higher biomass yield and numerical cell yield at a reduced growth rate^6^. This was the first experimental illustration of (cell) yield selection.

We exploited this emulsion protocol to evolve *E. coli* MG1655. Unexpectedly we found no reduction of overflow metabolism after experimental evolution, but rather a decrease in cell size, specifically during growth on the overflow metabolite acetate. This led to an increased numerical cell yield (number of cells/mole glucose). Our experiments and a mathematical model show that in a feast-famine regime fast growth and overflow metabolism followed by the consumption of the overflow metabolite can lead to a higher fitness compared to full respiration of the substrate.

## Results

### Selection for cell yield does not decrease overflow metabolism

The serial propagation of individual microbial cells in emulsion droplets resembles millions of parallel batch cultivations, each inoculated by a single cell. In each droplet, cells can grow for a limited number of generations (in our case 5 to 6, set by the average droplet size and the available carbon source) before they reach stationary phase. After such a growth period, all droplets are merged, the cells are mixed and diluted, and used to inoculate new droplets for the next round of growth. This protocol (figure 1B) selects for increased numerical cell yield per mole glucose. We wondered if this selection protocol would lead to *E. coli* strains with an increased metabolic efficiency (i.e. yield of gram biomass/mole glucose) and reduced overflow metabolism (more respiratory). To investigate this, we propagated *E. coli* MG1655 for 53 transfers in emulsions and determined the growth characteristics of 90 isolates (Supplementary Figure S1). One of these isolates, designated IR1, was characterized in more detail as it showed an increased maximal optical density compared to the wild type.

Besides the increase in four biomass measures (dry weight, total cell volume, total protein content, optical density) of 9-13%, strain IR1 also showed an increased growth rate on glucose (39% higher) (Fig. 1C-F). Genome sequencing (Supplementary Table S3) revealed an 82bp deletion in the *rph/pyrE* region that was previously characterized^2,9^. This mutation alleviates a documented pyrimidine production deficiency of the ancestral strain^10–12^ that limits growth on minimal medium, but not on rich medium. Growth experiments confirmed that IR1 also displays this phenotype (Supplementary Figure S3). We also found that during the glucose-growth phase, IR1 shows a 98% decrease in the production of pyrimidine intermediates orotate and dihydroorotate (Supplementary Figure S2). Together this data indicated that the increase in yield and rate is partially due to mutations that relieve the pyrimidine deficiency of the ancestral strain.

Mutants that showed a shift from fermentation towards respiration were not found. The amount of acetate produced per mole of consumed glucose was not altered significantly in the evolved mutant IR1 (Figure 1G). In an additional attempt to isolate mutants with altered overflow metabolism activity, we propagated strain IR1 for another 25 transfers in emulsion droplets. The subsequent characterization of 15 evolved strains showed that all strains still produced acetate (see Supplementary Table S2).

Another phenotypic change of the isolated mutant IR, was a decrease in cell volume of 11% compared to the ancestral strain. This cell size decrease was especially obvious during the second growth phase on acetate, and it corresponds to an increase in cell number of 23% (Figure 1C) when the glucose is depleted. Re-sequencing revealed a mutation that leads to a stop codon in the *ygeR* gene (Supplementary Figure S2). This gene is involved in septum formation and cell division, and deletion of it has been shown to reduce cell length^13^. The fact that we isolated mutant strains with reduced cell size and that we were not able to identify strains with decreased overflow metabolism led to an alternative hypothesis: In a feast-famine environment where the evolutionary pressure is on the overall *numerical* cell yield, overflow metabolism is actually more efficient than complete respiration when the consumption of the overflow product acetate is taken into the equation.

### Biphasic substrate utilization optimizes fitness in feast-famine environments

During growth in batch culture until complete exhaustion of all the carbon sources, microbes are subjected to continuously changing conditions. This resembles a feast and famine cycle with two feast phases (a first phase on glucose and a second phase on acetate), followed by carbon source exhaustion/ famine phase. If the carbon source is a fast fermentable sugar such as glucose then the first phase will be characterized by a high growth rate and a relatively large cell size^14^. As long as the sugar concentration is high, maximizing the growth rate leads to the highest number of offspring produced per unit time. At this stage it is not relevant if the cell is metabolically inefficient, as there is sufficient substrate available. However, when the initial “fast” substrate becomes limiting, cells will switch to growth on the produced overflow metabolite, acetate in the case of *E. coli*. Growth on an overflow metabolite is always slower than on the initial fast substrate and slow growth is correlated with smaller cell sizes^14–18^.

We show that already during the onset of the second growth phase, the cells started to reduce in size to eventually reach a cell volume in stationary phase that is less than half of what it was during exponential growth on glucose (Figure 1E). During this cell size adaptation period, making new offspring cells costs less nutrients than during steady-state growth on acetate, because a mother cell is now larger than a daughter cell and likely does not need to double in size before division. Immediately after glucose depletion, cells therefore need to produce proportionally less biomass to divide into two daughter cells. This leads to a proportionally higher increase in cell number compared to the increase in biomass during the transition from glucose to acetate (Figure 1C-D). This reduction in cell size during the acetate growth phase is substantially larger compared to conditions where no biphasic growth was possible (see Supplementary Information 1). Together, the large differences in cell size on glucose and acetate, the identification of mutants with a smaller cells size and the consequences of this phenotype on the numerical cell yield suggest significant fitness effects of cell size in feast-famine conditions.

### Rate and cell yield selection effects of biphasic carbon source utilisation

The widespread interpretation of the evolutionary benefit of overflow metabolism is biased towards the rate selection argument. However, repeated cycles of growth with finite nutrient supply broadly occurs in nature, and it is likely to have played an important role during the evolution of metabolic strategies. To address whether biphasic substrate utilization is an evolutionary advantageous strategy to maximise the number of offspring in environments with finite resources, we made a mathematical model (see Supplementary Information 2 for derivation and description). The model calculates the number of cells that are made from an initial amount of glucose, given the known biomass yields on glucose and acetate, the fraction *ϕ* of glucose converted into acetate, the cell sizes on glucose and acetate, and the growth rates on glucose and acetate. In agreement with experiments, the growth rate on glucose is a function of *ϕ*: during pure respiratory growth (at *ϕ* = 0) the growth rate is lower than during overflow metabolism (intermediate *ϕ*) and when only acetate would be made from glucose (*ϕ* = 1), the growth rate on glucose is zero. A schematic of the model is shown in Figure 2A-B. We used the model to determine the fitness in the case of selection under feast and famine cycles (see Supplementary Information 2).

**Figure 2.**
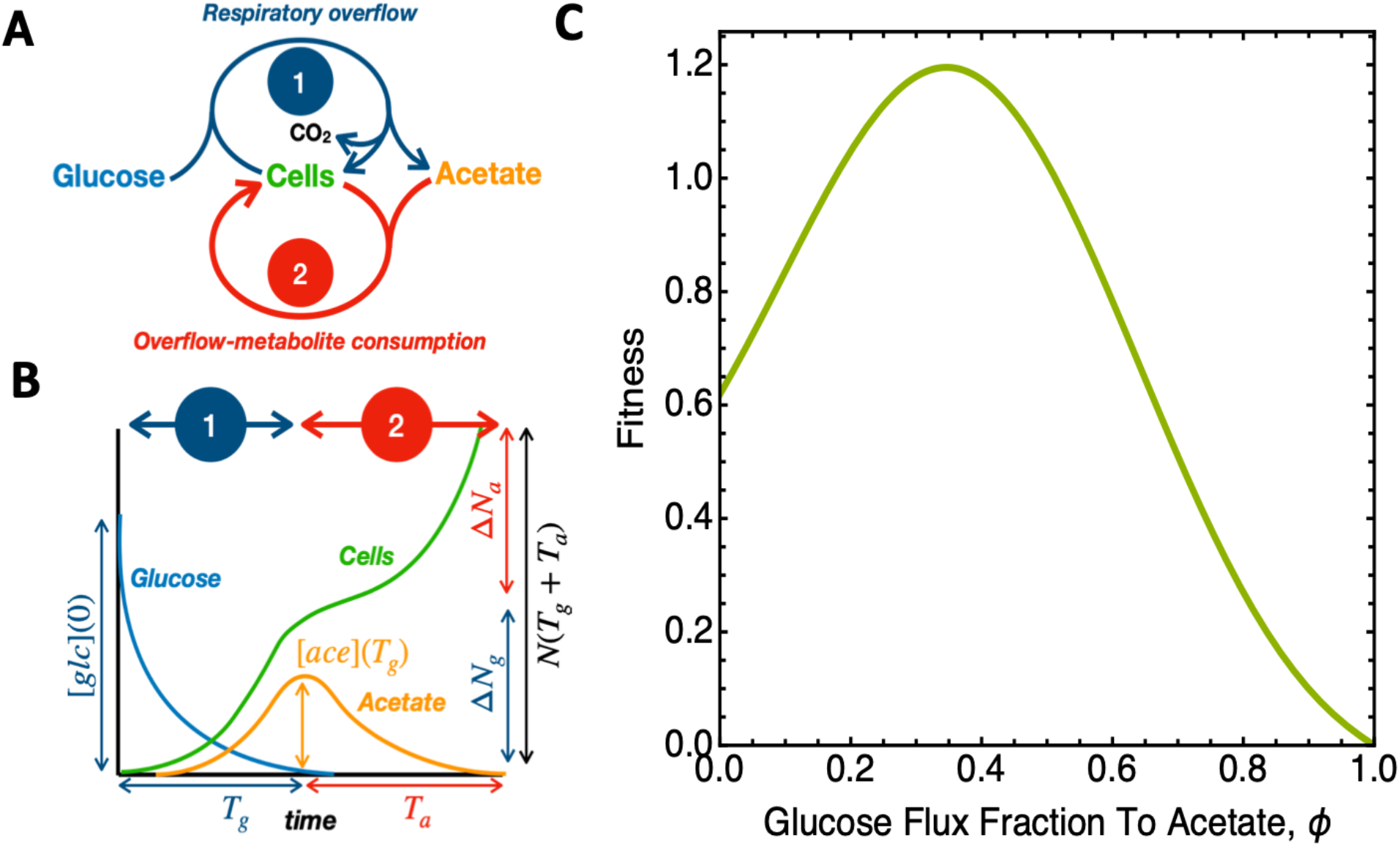
Modelling the fitness consequences of biphasic growth for Escherichia coli. **A.** Schematic overview of metabolic processes associated with biphasic growth: glucose consumption, acetate formation and growth during the first phase and acetate consumption and growth during the second phase. **B.** An illustration of the concentrations and cell abundance as function of time during biphasic growth and introduction of terminology. **C.** We used the model to calculate the fitness (F, see Supplementary Information 2) as function of the fraction of the glucose flux directed to acetate formation. Fitness is defined as the logarithm of the factor increase in the number of cells divided by the time until carbon source depletion, i.e. 1/(Tg+Ta) ln N(Ta+Tg)/N(0), in agreement with classical evolutionary biology. Parameters values can be found in the table of Supplementary Information 2.

Figure 2C shows that the qualitative model predicts maximal fitness not when cells fully respire (*ϕ* = 0) but rather when a fraction of the consumed carbon is first converted to acetate. A fully respiratory cell has a lower fitness than this strategy, indicating that selection during feast-famine conditions selects for a phenotype that combines fast growth (overflow metabolism on glucose) with small cell/high cell number (growth on acetate).

## Discussion

Studies describing overflow metabolism as inefficient typically do not include the consumption of overflow metabolites when comparing it to yields of fully respiratory cells. This reasoning is valid during constant conditions of high glucose concentrations where overflow metabolism indeed reduces the maximum attainable ATP (and gram biomass) yield of the utilized carbon source^5,19–21^. It does however not apply when the carbon source is depleted, e.g. in batch cultures when utilization of a carbon source is followed by the consumption of the overflow metabolites. Therefore, for a fair comparison, the additional biomass yield on the overflow metabolites needs to be considered as well. In this case it is not immediately clear whether fully respiratory growth is a better evolutionary strategy than overflow metabolism and biphasic growth.

Another important point is that natural selection acts on the number of viable offspring rather than the amount of produced biomass. Interestingly, many bacteria become larger when they grow faster; also *E. coli* is known to have a positive correlation between growth rate and cell size^15–18^. This occurs when the generation time is shorter than the DNA replication time, necessitating mother cells to start DNA replication for their future progeny^22–24^. During evolution on a finite nutrient supply however, it pays off to make a higher number of cells per unit nutrient, and this can be achieved by making smaller cells. A fast growth rate that supports an evolutionary advantage under nutrient excess, therefore has a trade-off with a lower number of offspring per unit of substrate. This trade-off is driven by cell size and independent from metabolic efficiency. An optimization problem thus occurs, where small (but slow) phenotypes would generally be outgrown by faster growing cells during nutrient competition. We circumvented this trade-off by serially propagating cells in emulsion droplets where single cells are allowed to grow until nutrient exhaustion in physically separated environments. In this regime mutants that produce more offspring will seed more droplets in the next round and increase in frequency.

Against initial expectations, full respiration was not found to be a competitive strategy when rate selection was excluded. While selection in emulsion led to increased metabolic efficiency, biomass- and cell-yield in *L. lactis*^6^, we found that in *E. coli* the utilization of overflow metabolism was unaffected. However, we did select for a mutant with a smaller cell size, which is possibly linked to a single base deletion in *ygeR, a* gene known to affect the cell size^13^. We identified two reasons for the difference in the outcome between the *L. lactis* study and our experiments i) *E. coli* is able to consume the overflow metabolite which *L. lactis* cannot, and ii) when growing on the overflow metabolite, the cell number of *E. coli* increases disproportionately compared to the biomass increase.

Our results show that overflow metabolite production and its subsequent utilization maximizes the number of produced offspring irrespective of the available substrate concentration. At high substrate concentrations the growth rate will be maximized while at limiting substrate concentrations a minimization of cell size (maximization of cell number) will occur after the initial “fast” substrate is depleted. This allows the optimization of fitness in dynamic environments. Maximizing the production rate of offspring through sequential substrate utilization seems a very elegant solution that might apply to numerous organisms. For instance the secretion of ethanol, lactate or acetate and their subsequent consumption are broadly encountered^25–27^. Like for *E. coli*, other organisms showing such behaviour are also described to have a positive correlation between growth rate and cell size^6,18,28^. This makes it plausible that a similar fitness maximization strategy in dynamic environments might apply to a broad range of organisms.

Attempts to engineer Crabtree negative yeasts and *E. coli* have been only partially successful and typically the obtained strains are growing poorly or showing unexpected metabolic activities^29^. This points to overflow metabolism being a robust property of the metabolic networks that are difficult to overcome. However, Crabtree negative yeasts are known and it would be interesting to understand under which conditions they might have evolved^30,31^.

There are numerous computational approaches that predict the shift from respiration towards overflow metabolism, based on biochemical and biophysical constraints. These studies include for example flux balance analysis (FBA), where the optimal metabolic flux distribution is predicted that supports high growth rates; trade-offs due to membrane crowding; and optimization of proteome allocation to minimize the investment associated with the costly respiratory machinery^32–38^. However, besides some studies in yeast where the main argument is that overflow metabolites lead to a fitness advantage due to their toxicity for competing organisms (Make-Accumulate-Consume hypothesis)^27,30,39^ the role of evolutionary perspectives are less investigated. The discussion of overflow metabolism in literature and textbooks typically neglects the fact that the secreted overflow metabolite is often further metabolized. Our results argue for the consideration of the consumption of the overflow metabolite as it likely played a role during evolution in natural environments. This is corroborated by a recent study on the regulation of overflow metabolism^40^. Full respiration and reaching the maximal biomass yield are only seen at low substrate concentrations. This might be a rather artificial situation which is created for instance in chemostats. In nature cells are often exposed to dynamic feast-famine cycles, and spatial structure that leads to variations in the selection pressure.

If rate selection would be the only force to favour overflow metabolism one might expect higher acetate fractions to be produced by the wild type strain. The fact that in most cases only a small fraction of the flux is directed towards acetate argues for other (additional) selective forces, such as maximizing cell number. In studies by LaCroix et al.^2^ and Long et al.^38^ rate selection was applied to the same *E. coli* MG1655 wild type strain. They found a positive correlation between the acetate production and growth rates in the wild type and evolved strains. Such an increase in overflow metabolism at increasing growth rates has been reported in a number of other studies as well^3,7,41,42^. This might indicate that the natural environment from which the wild type strain was isolated was not purely selective for growth rate.

Another aspect of natural environments is the possibility of multiple species or strains coexisting. A ‘cheater’ strain that consumes the acetate before the producer strain can benefit from it would potentially argue against a beneficial effect of overflow metabolism beyond growth rate maximization^43^. However, growth on acetate is typically slow, so even if such a strain would be present at high frequency in the population, its consumption rate would still be significantly lower than what is needed to fully deplete this substrate before the fast growing/acetate producing biomass starts consuming it. In order to pose significant competition, the acetate consumer would therefore need to grow at a similar rate as the acetate producer. Almost all growth rates we found for diverse organisms on acetate were substantially lower than this (see Supplementary Table S5 for a number of examples of glucose and acetate growth rate ratios), except for an *Acinetobacter* strain that displays an exceptionally high growth rate on acetate of 0.91 h^−1^ ^44^.

In conclusion, this study shows an unexplored consequence of overflow metabolism. It is not only a strategy for fast growth at excess substrate conditions, but it also maximizes the number of offspring in dynamic conditions with regular episodes of finite substrate concentrations. From an evolutionary perspective overflow metabolism is potentially selected for because it maximizes the number of offspring in such dynamic environments, be it through a fast growth rate or the minimization of the cell size. Our results argue for a revisited perspective when investigating metabolic strategies, as currently consumption of overflow metabolites and the role of cell numbers during selection are neglected when fermentation is compared to full respiration.

## Material and Methods

### Strains and media

For experimental evolution *Escherichia coli* K12 substr. MG1655 (isolate BOP27 obtained from Palsson lab at UCSD) was used. The experiments were initiated from a single colony glycerol stock which was cultured in 3 biological replicates. Strains were cultured at 37°C using M9 minimal medium^2^ supplemented with 2.5 mM glucose. The same medium was used throughout this study.

### Culturing in emulsion

A fully-grown cell culture of *E. coli* was diluted to a concentration of ~2.10^6^ cells.ml^−1^ in M9 medium supplemented with 2.5 mM glucose. Using 700 μl of the diluted cell suspension and 300 μl Novec HFE 7500 fluorinated oil (3 M, Maplewood, MN, USA) containing 0.1% Pico-Surf™ surfactant (Dolomite Microfluidics) emulsion where prepared as described earlier^6^. In such an emulsion approximately 1 in 10 droplets (~50 μm in diameter) will be inoculated with a single cell (following a Poisson distribution). After overnight incubation at 37°C, the emulsions were broken using perfluoroctanol (PFO)(Alfa Aesar) and the cultures were diluted in fresh medium to repeat the process^6^.

To ensure aerobic growth was supported, the oil was saturated with pressurized atmospheric air prior to use. Sufficient O_2_ supply was confirmed by measuring growth on a non-fermentable carbon source (data not shown).

### Growth curve measurements

Strains of interest were precultured and propagated to a 96-wells plate. The spaces between wells were filled with 0.9% saline solution, and the plate sealed with parafilm to reduce evaporation from the wells. Growth was measured every 5 minutes, with shaking in between, overnight in a SpectraMax® plate-reader, at 600 nm and 37°C. Using the R software environment, growth rates and maximum ODs were determined per strain as described earlier^6^.

Alternatively, growth was measured manually. Strains were grown in 250 ml Erlenmeyer flasks, shaking at 220 RPM, at 37°C. Every 30 minutes samples were taken to measure OD_600_ in cuvettes.

### Initial screening after evolution

After the experimental evolution 90 single colonies were randomly picked from the three parallel cell cultures and transferred to a 96-wells plate containing 200 μl M9 + 2.5mM glucose per well. Six wells were inoculated with the wild type strain. After overnight growth, 20 μl from each well was transferred to a 96-wells plates containing fresh medium for proper growth curve measurements. Glycerol was added to a final concentration of 12% and the plates were frozen at - 80°C. Growth curves were analysed for growth rates and maximal ODs. Several strains per replicate culture were selected for further analysis, based on the maximal OD compared to the wildtype.

### Cell number and volume measurements

Cell number and volume were determined using the Coulter Counter® Multisizer 3 (settings: aperture 30 μm; kd 41; current 800 μA; gain 16; sizing threshold 0.3 μm) after adding 50 μl cell culture to 10 ml ISOTON II electrolyte.

### Metabolic product measurements with HPLC

HPLC analysis was done on the (0.2 μm) filtrate of the culture supernatant to determine the metabolic products. The HPLC (Shimadzu LC20, 300×7.8mm) with an ion exclusion column Rezex ROA Organic Acid H+ (8%), was run with 5 mM H_2_SO_4_, kept at 55°C at a flow rate of 0.5 ml.min^−1^. Calibrations were done with formate, lactate, acetate, orotate and dihydroorotate.

### Glucose determinations

Given the high concentration of phosphate in the M9 medium, and the overlap of this compound with glucose in the HPLC chromatogram, enzymatic glucose determinations were used to complete the metabolic profile. This was done by preparing a reaction mixture containing (final concentrations) PIPES pH=7.0 (14.627 mM), NADP^+^ (0.286 mM), ATP (0.571 mM), MgSO4 (1.428 mM), hexokinase (30 U/ml), and glucose-6-phosphate dehydrogenase (35 U/ml). In a 96-wells plate, 22 μl of sample or standard containing known concentrations of glucose were added to each well, then 178 μl of reaction mixture was added to every sample/standard. This was mixed, incubated for ~30 minutes at 30°C, and measured at 340nm until the curves were stationary. Glucose concentrations were calculated using the calibration curve (measured in duplicate), based on the final A_340_.

### Protein content, dry weight

Total protein content of the selected strains was determined following the protocol of the Pierce® BCA Protein Assay Kit. To prepare the samples for this assay, 2 ml of culture was transferred to an Eppendorf tube, washed in 0.9% saline solution, and the cell pellet was re-dissolved in 200 μl of saline. 50 μl of SDS (10%) was added, and the samples were incubated at 90°C for one hour. Of these samples, and a set of calibration samples with known BSA concentrations, 25 μl was transferred to a 96-wells plate and 200 μl of reagent (Pierce® BCA Protein Assay Kit; Reagent A: Reagent B = 50:1) was added. Sample and reagent were mixed by shaking on a plate-shaker for 30 seconds, then incubated at 37°C for 30 minutes. Protein content was subsequently determined in a SpectraMax® plate-reader at 562 nm, and data analysed using the BSA-calibration curves (measured in duplicate), based on the final A_562_.

The dry weights of the selected strains were determined by filtering 50 ml of cell culture through a previously dried 0.2 μm pore nitrocellulose filter using a vacuum pump, drying these filters at 60°C to a constant weight and determining the difference in weight before and after the application of the samples to the filters, using a precision scale.

### Genome re-sequencing

Genomic DNA was isolated using a Promega Wizard DNA purification kit. The quality of DNA was assessed with UV absorbance ratios by using a NanoDrop apparatus. DNA was quantified by using a Qubit dsDNA high-sensitivity assay. Paired-end resequencing libraries were generated using an Illumina Nextera XT kit with 1 ng of input DNA total. Sequences were obtained using an Illumina Miseq with a PE500v2 kit. The Breseq pipeline^45^ version 0.23 with bowtie2 was used to map sequencing reads and identify mutations relative to the E. coli K-12 substr. MG1655 genome (NCBI accession NC_000913.2). All samples had an average mapped coverage of at least 25.

## Acknowledgements

We would like to thank Vera Benavente, Tim van Wagensveld and Wouter W. Woud for technical assistance. IR and FJB were financed by NWO-VIDI project 864.11.011.

## Author contribution

IR, BT and HB conceived the study. IR carried out the experiments. AF re-sequenced evolved strains. HB supervised the study. FJB and WG constructed mathematical models. IR, FJB and HB analysed the data and wrote the paper. All authors read and commented on the manuscript.

## Supplementary Information

### Supplementary Information 1. Comparison of cell size under different growth conditions

An underlying assumption in the hypothesis above is that the cell size after entering stationary phase when growing on acetate is smaller than when entering stationary phase after growing on glucose. We found that when wildtype cells are growing in a glucose limited batch culture where the final divisions are on the overflow metabolite acetate, the cell size in stationary phase is approx. 55% smaller than the cell size during exponential growth on glucose. As strain MG1655 always produces acetate in a batch culture on glucose investigating the entering of stationary phase without acetate exposure in batch culture is not possible. To mimic the effects on cell size when going into stationary phase at a faster growth rate than on acetate we prepared nitrogen limited batch cultures where no biphasic growth is observed. The results showed that the final cell size after going into stationary phase with nitrogen limitation is approx. 24% smaller than during exponential growth (see Supplementary Figure 4), which is significantly bigger than cells entering stationary phase after growth on acetate.

A second assumption is that the growth rate reduction that comes with full respiration does not lead to a cell size decrease that combined with its effect on increasing biomass yield would lead to a higher cell yield than growth on acetate. We found 5 studies with *E. coli* where maximum growth rates and growth rates that still allow full respiration are reported (see Supplementary Table 4). All five show that a growth rate reduction to 77% - 59% of the maximum growth rate is sufficient for a strain to change metabolism towards full respiration. In our case the balanced growth rate on acetate is 28% and 31% of the maximum growth rate for MG1655 and IR1 respectively. This is therefore well below the growth rate reduction required to allow full respiration suggesting that the growth rate reduction on acetate as a substrate adds to the cell number through making smaller cells.

### Supplementary Information 2. Model description

#### Introduction

The aim of this model is to evaluate the fitness of biphasic metabolism, incl. overflow metabolism, and compare it to the fitness of a pure-respiration strategy; to identify which parameters determine the winning metabolic strategy, i.e. the one favoured by evolution when cells are confronted with a finite amount of sugar that they can consume until its depletion.

We assume that the cells deplete a finite amount of sugar. This sugar amount limits the final number of produced cells, as all other nutrients are assumed in excess. Selection of cell number yield takes place when single cells are allowed to grow, while they are physically separated in emulsion droplets containing glucose-limited medium. In this scenario the total time of growth does not matter. Selection of growth rate occurs in a well-mixed environment in a batch culture, also until all carbon sources have been depleted. In both cases, the winning evolutionary strategy made the most offspring after all carbon sources, i.e. the limiting nutrients, have been depleted (sugar and overflow metabolites).

#### Model equations

The number of cells made from a finite starting amount of glucose, *glc[0]*, when a fraction *ϕ* of the glucose uptake rate goes to acetate formation equals,

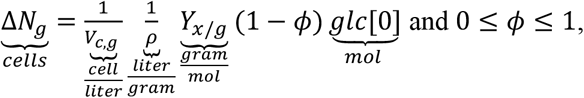

with the units of *each factor* explicitly indicated: *V_c,g_*= the volume of a single cell during growth on glucose, *ρ*= the mass density, and *Y*_*x/g*_= the gram biomass yield on glucose. Note that no cells are made when all glucose, during the glucose phase, is converted into acetate (i.e. *ϕ* = 1). After the glucose growth period, with duration *T*_*g*_, during which Δ*N*_*g*_ cells were made, the total number of cells equals,

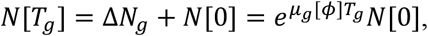

with *N*[0] as the starting number of cells, such that

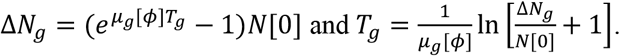

The function *μ*_*g*_[*ϕ*] denotes the *growth rate during the glucose phase* and is dependent on the fraction *ϕ* of glucose converted into acetate. Chemostat experiments indicate that fully respiring cells, with *ϕ* = 0, grow slower than partially-overflowing cells with *ϕ* ≈ 0.2. And, cells that only overflow, i.e. *ϕ* = 1, clearly do not grow on glucose at all. The dependency of the growth rate on glucose on *ϕ*, i.e. *μ*_*g*_[*ϕ*], therefore has to obey the following experimental observations:

1. when *ϕ* = 0, it equals the growth rate at pure respiration, *μ*_*g*_[0] = *α μ*_*a*_, which exceeds *μ*_*a*_ (the growth rate on acetate); thus, *α* > 1, as a fully respiring cell growing on glucose grows faster than a cell growing on acetate (given chemostat and batch data).
2. at *ϕ* = 1, the growth rate on glucose equals 0 as all glucose is converted into acetate.
3. a *ϕ*-value *ϕ*_*peak*_ exists, 0 < *ϕ*_*peak*_ < 1, with a growth rate *μ*_*g*_(*ϕ*_*peak*_) that exceeds the growth rate at pure respiration *α μ*_*a*_ (and *μ*_*a*_).

Thus, the function *μ*_*g*_[*ϕ*] rises from a nonzero starting value *μ*_*g*_[0] > *μ*_*a*_ > 0 to a maximal value at *μ*_*g*_[*ϕ*_*peak*_] and then decreases again to 0 at *μ*_*g*_[1]. This behavior is an extrapolation of experimental data. The following function for the growth rate on glucose suits our purposes (with the Greek letters denoted positive parameters),

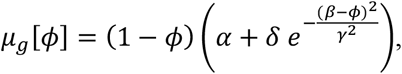

which equals zero at *ϕ* = 1 and is positive at *ϕ* = 0 where its value equals *αμ*_*a*_. Note that this function shows the desired behavior, but is otherwise arbitrary.

The amount of acetate produced during the glucose growth period equals,

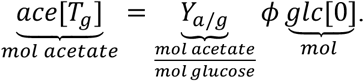

The number of cells that we can make from this amount of acetate equals,

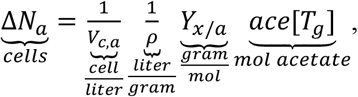

with *V*_*c,a*_ as the volume of a cell during growth on acetate and the *Y*_*x*/*a*_ as the gram biomass yield on acetate. (Note that we neglect that from glucose-grown cells, more cells can be made than expected from the acetate yield and acetate amount, because glucose-grown cells are larger than acetate cells (see main text). This effect we neglect in the model, to keep it simple.)

Thus, the number of cells at the end of the experiments, after glucose and acetate growth, equals

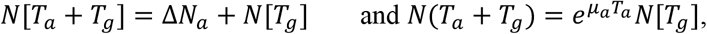

with T_*a*_ as the duration of the acetate growth period and *μ*_*a*_ as the growth rate on acetate. These last two equations allow us to determine the growth period on acetate,

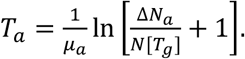

The total number of cells made from the starting amount of glucose equals,

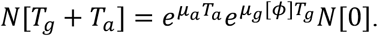

The total number of cells made equals Δ*N* = *N*[*T*_*g*_ + *T*_*a*_] — *N*[0].

We define the growth factor as

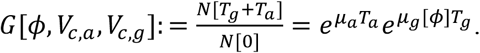

and the geometric fitness as

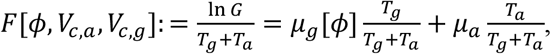

which equals the time-averaged growth rate, since 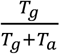 equals the fraction of time spent in phase 1 (glucose phase) and 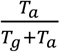 equals the fraction of time spent in the acetate phase. The *F*[*ϕ*, *V*_*c,a*_, *V*_*c,g*_] is plotted in Figure 2C.

#### A complication: cell size and growth rate are related for *E. coli*

*E. coli* has a fixed DNA replication time. This means that at long generation times (low growth rates) enough time exists between cell birth and division to replicate a single copy of DNA. This is not the case when the generation time is shorter than the DNA replication time: then DNA replication occurs continuously throughout the cell cycle and cells have started replication of DNA copies for future progeny.

As a result, cells that grow fast have multiple origins of replication. It has been shown that the exponential relation between cell size and growth rate can be understood to result from the fact that *E. coli* maintains a constant ratio of cell size over the number of origins of replication. What matters for our model is that wild type cells obey the following relation: *V*(*μ*) = *V*(0)*e*^*εμ*^with *V*(0) = 0.28 *μm^3^* and *γ* = 1.33 *hr*^1^.

**Table.** The used parameters for Figure 2.

| Name | Value |
| --- | --- |
| Cell density | $280 \times 10^{-15} \text{ g}/\mu\text{m}^3$ |
| Biomass yield on glucose | 88 g/mol |
| Biomass yield on acetate | 20 g/mol |
| Acetate yield on glucose | 2 mol/mol |
| Cell volume on glucose ($V_{c,g}$) | $1 \mu\text{m}^3$ |
| Cell volume on acetate ($V_{c,a}$) | $0.5 \mu\text{m}^3$ |
| Growth rate on glucose ($\mu_g$) | $1 \text{ hr}^{-1}$ |
| Growth rate on acetate ($\mu_a$) | $0.3 \text{ hr}^{-1}$ |
| $\alpha$ | 0.2 |
| $\beta$ | 0.5 |
| $\gamma$ | 0.4 |
| $\delta$ | 2 |

## Supplementary Figures

**Supplementary Figure 1:**
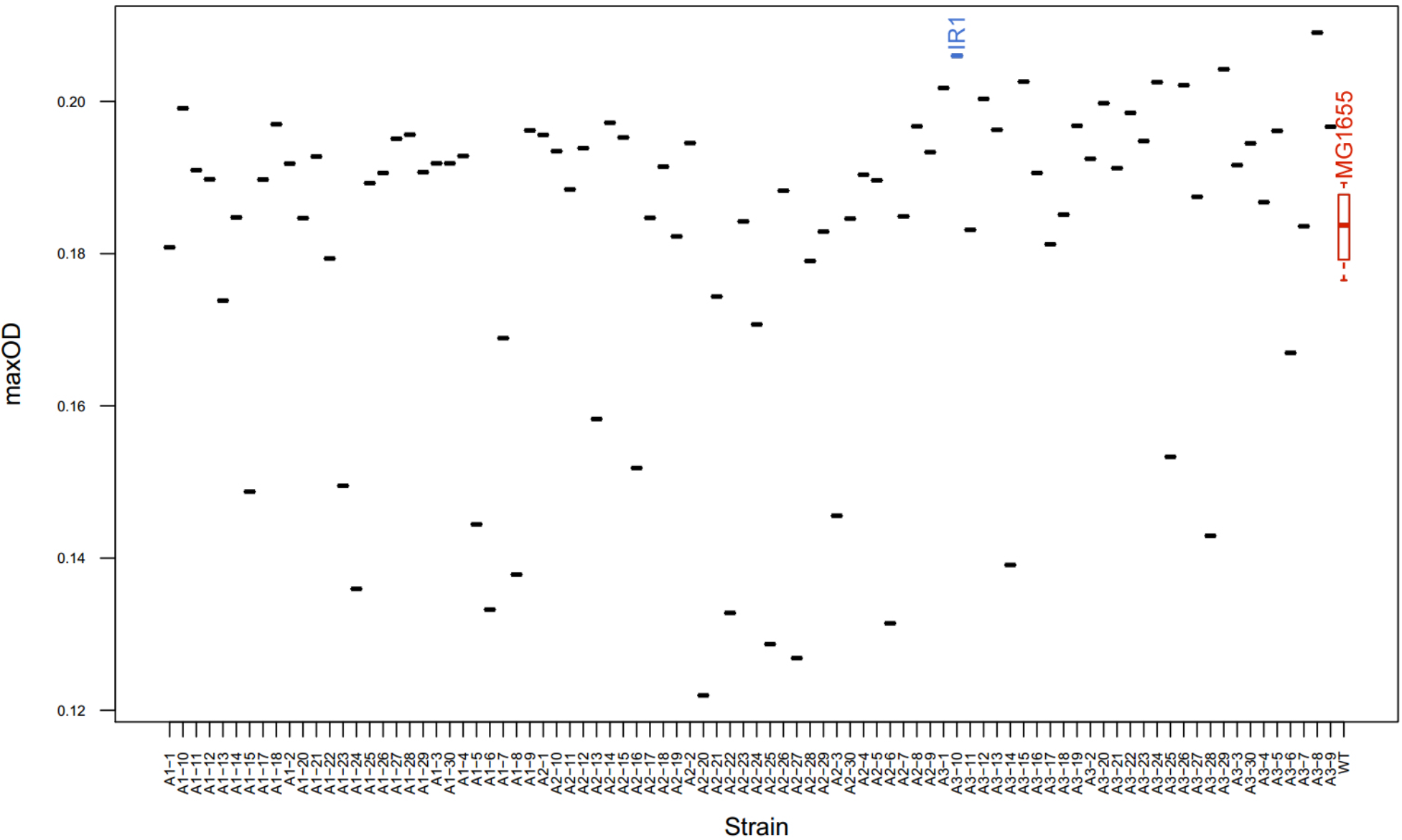
After 53 propagations in emulsion, three replicate populations A1, A2 and A3 were plated, and 90 single colonies were picked. The maximal optical densities at 600nm of these evolved strains were compared to that of the wildtype (boxplot of 6 replicates), to screen for strains with a suspected increase in yield. Strain IR1 was selected for extensive characterization based on the initial screening measurements.

**Supplementary Figure 2:**
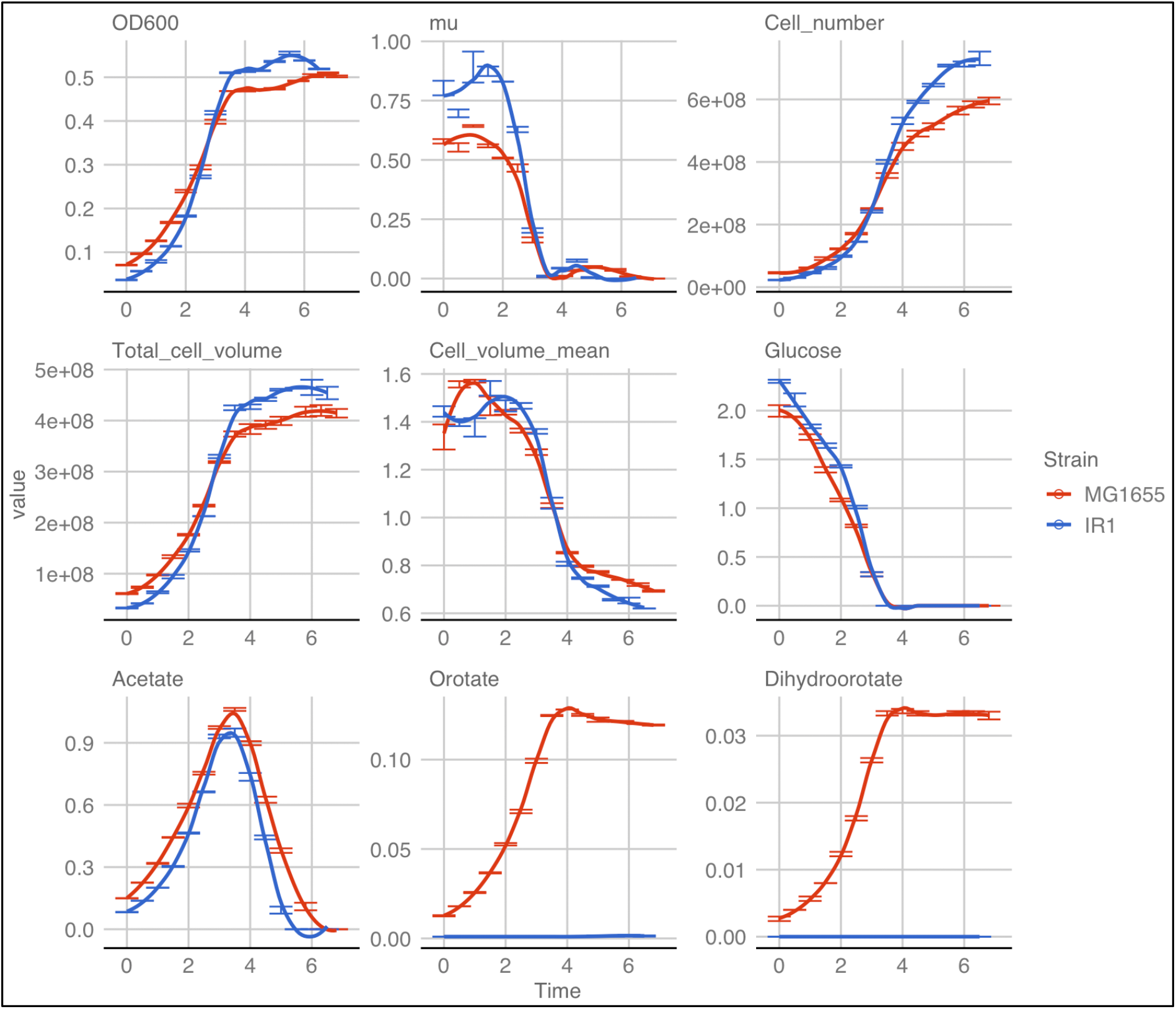
Growth characteristics from batch cultures. Units: Growth rate mu [h^−1^], Cell_number [number/ml culture], Total_cell_volume [μm^3^/ml culture], Cell_volume_mean [μm^3^], Glucose [mM], Acetate [mM], Orotate [mM], Dihydroorotate [mM]. Error bars are standard errors of the mean, n=3. For completenes the panels from Fig 1 C-E (main text) are shown here as well.

**Supplementary Figure 3.**
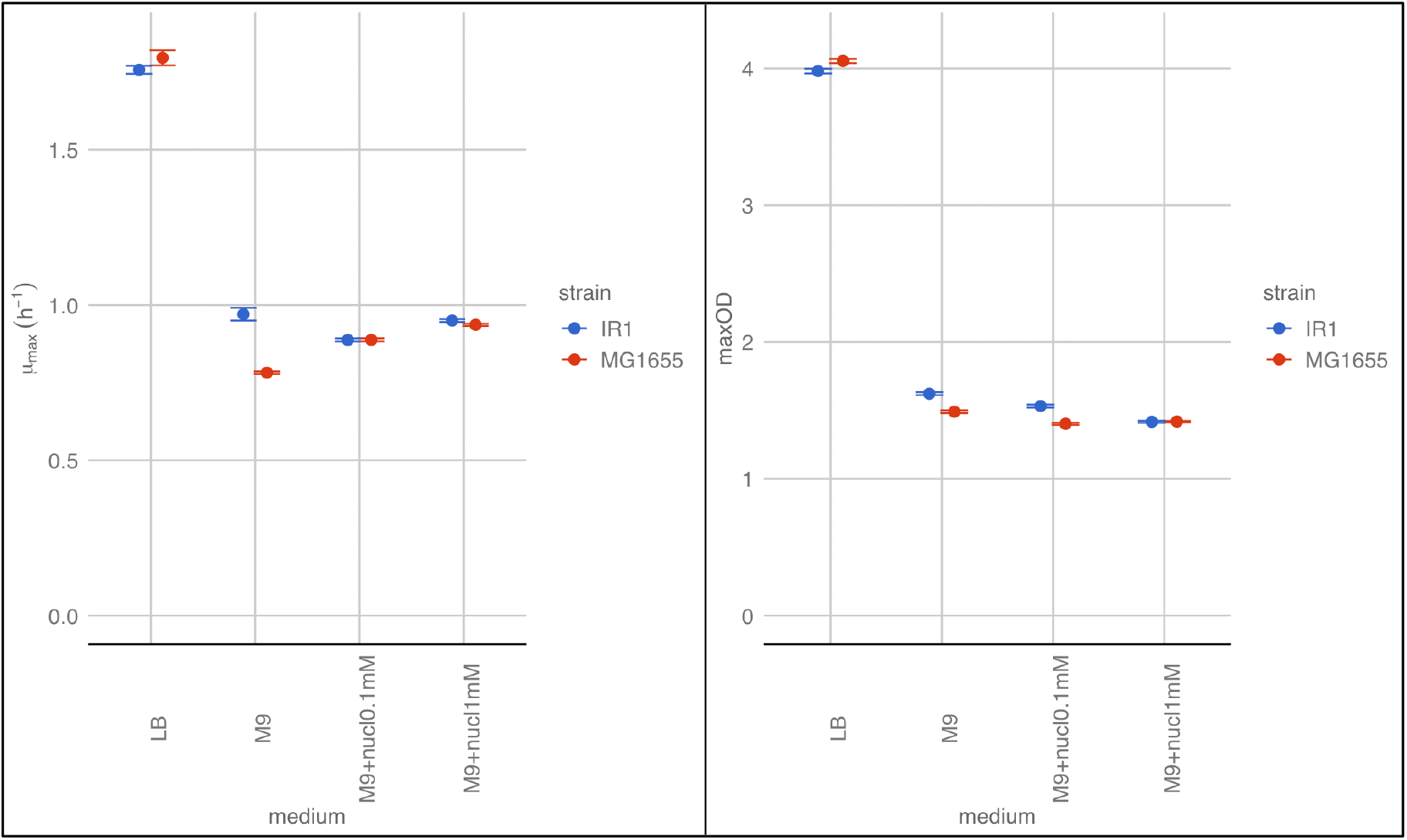
Phenotypic properties of the wild type strain and IR1. Phenotypes associated with a mutation in the rph-pyrE region that strain IR1 acquired when evolved on M9 minimal medium, is consistent with an earlier described medium adaptation^2–5^. On rich LB medium the ancestral wildtype strain has a higher maximal specific growth rate (μ_max_) and final optical density (maxOD) than strain IR1. When grown on M9 medium however, the pyrimidine production deficiency of the wildtype strain leads to a reduced μ_max_ and maxOD. If the minimal medium is supplemented with free nucleotides (0.1 or 1 mM), this disadvantage of the wildtype strain is alleviated. Means with SEM are shown, n=6.

**Supplementary Figure 4:**
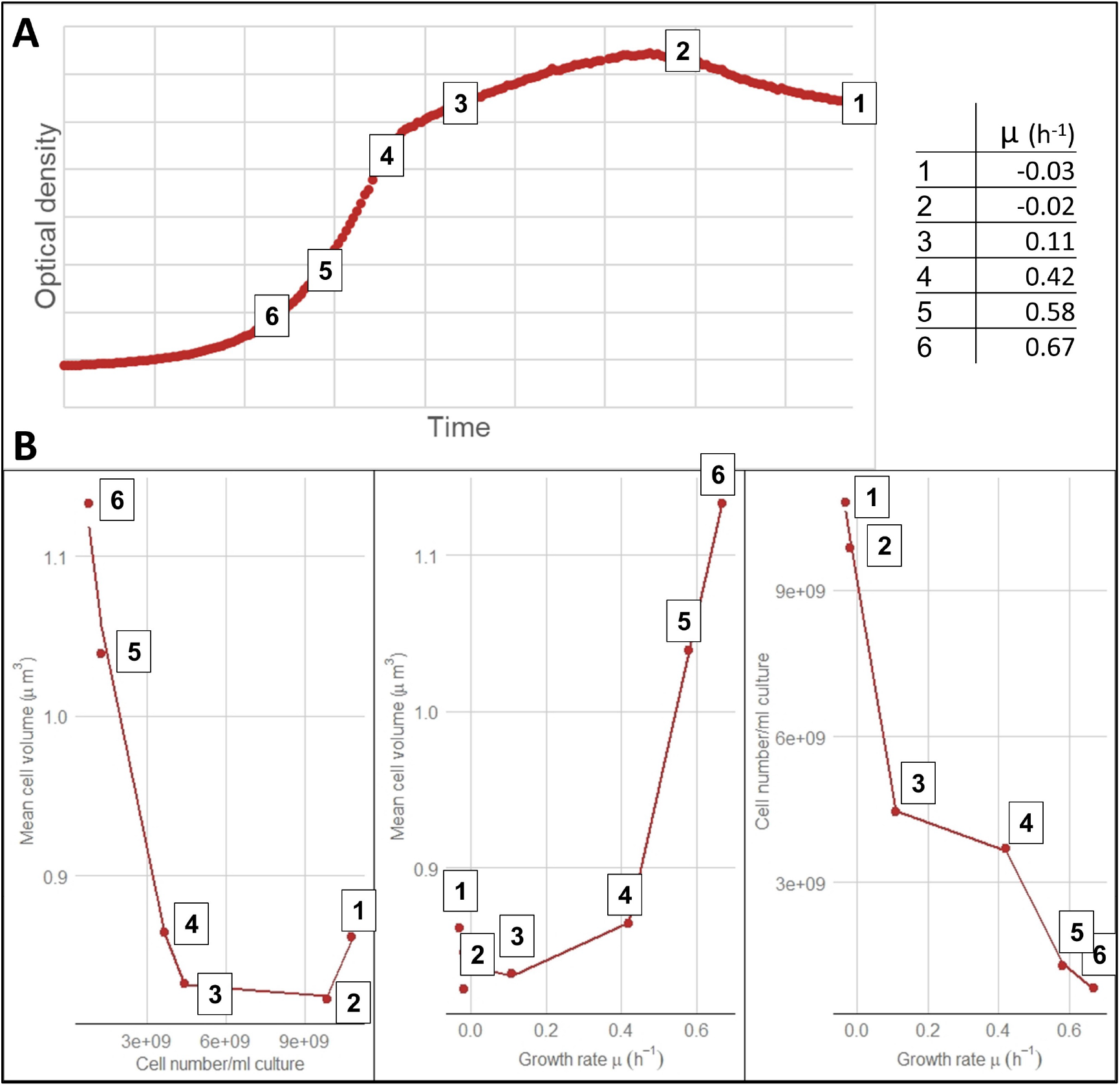
**A.** Wild type MG1655 was grown in batch on M9 supplemented with 50 mM glucose. In this case the growth medium is nitrogen limited instead of carbon limited hence cells still grow on glucose when going into stationary phase (no growth on acetate). Using a dilution series of different inoculation densities samples were taken at various timepoints throughout the growth curve for Coulter Counter measurements (i.e. cell volume and cell number). The growth rate (slope of log(OD)) was determined for different timepoints from mid-exponential to prolonged stationary phase. **B.** As the growth rate decreases, the cell volume also decreases while the cell number increases. In this nitrogen limited experiment, the decrease in cell volume from mid-exponential until stationary phase is around 24%. This decrease is significantly less than a decrease in cell volume of 55% observed for cells that become stationary in a glucose limited culture where a second growth phase on acetate occurs - see Supplemental Figure 2). Data was fitted with a smooth spline (smoothing parameter 0.3), n=2.

## Supplementary Tables

**Supplementary Table 1:** Overview of the phenotypic characterization from wild type MG1655 and evolved strain IR1. Mean and standard deviation are given (n=3). Growth rates and maximal ODs were measured in shake flask (SF), as well as microplate (MP). All other measures come from the shake flask cultures. The second growth phase on acetate is too short to reach balanced growth, and sample handling for shake flasks can lead to some temperature reduction, which can cause a reduction in apparent growth rates. The growth rate difference between the wild type and evolved strain were therefore confirmed using microplate growth curves to ensure balanced growth conditions.

|  | MG1655 |  | IR1 |  | IR1 %<br>of<br>MG1655 | t-test (p-value) |
| --- | --- | --- | --- | --- | --- | --- |
|  | mean | st.dev. | mean | st.dev. |  |  |
| cell number/ml culture | 5.95E+08 | 1.93E+07 | 7.31E+08 | 3.77E+07 | 123 | 0.0117 |
| glc exp cell volume ($\mu\text{m}^3$) | 1.57 | 0.01 | 1.54 | 0.05 | 98 | 0.4449 |
| stationary cell volume ($\mu\text{m}^3$) | 0.70 | 0.01 | 0.62 | 0.00 | 89 | 0.0002 |
| total cell volume/ml culture | 4.14E+08 | 1.44E+07 | 4.54E+08 | 2.12E+07 | 110 | 0.0615 |
| dry weight (mg/ml culture) | 0.163 | 0.006 | 0.181 | 0.006 | 111 | 0.0208 |
| total protein content ($\mu\text{g/ml}$ culture) | 306.0 | 8.5 | 345.4 | 10.2 | 113 | 0.0000 |
| maximal OD<sub>600</sub> - SF | 0.510 | 0.002 | 0.555 | 0.006 | 109 | 0.0027 |
| maximal specific growth rate<br>glc ($\mu_{\text{max}}$ ($\text{h}^{-1}$)) - SF | 0.579 | 0.010 | 0.805 | 0.008 | 139 | 0.0000 |
| maximal specific growth rate<br>acet ($\mu_{\text{max}}$ ($\text{h}^{-1}$)) during 2nd<br>growth phase - SF | 0.034 | 0.001 | 0.056 | 0.008 | 166 | 0.0359 |
| maximal OD<sub>600</sub> - MP | 0.531 | 0.013 | 0.570 | 0.008 | 107 | 0.0154 |
| maximal specific growth rate<br>glc ($\mu_{\text{max}}$ ($\text{h}^{-1}$)) - MP | 0.839 | 0.015 | 1.100 | 0.005 | 131 | 0.0003 |
| maximal specific growth rate<br>acet ($\mu_{\text{max}}$ ($\text{h}^{-1}$)) balanced<br>growth - MP | 0.224 | 0.007 | 0.294 | 0.011 | 131 | 0.0017 |
| fraction of C to acetate (end of<br>1st growth phase) | 0.159 | 0.005 | 0.150 | 0.003 | 94 | 0.0690 |
| fraction of C to pyrimidine<br>intermediates (end of 2nd<br>growth phase) | 0.051 | 0.001 | 0.000 | 0.000 | 1 | 0.0000 |

**Supplementary Table 2:** Strain IR1 was propagated for 25 additional cycles in emulsion, and 15 strains (3 per replicate evolution culture) were screened for acetate production after 3, 4 or 5 hours of batch growth. All strains still produced considerable amounts of acetate indicating that overflow metabolism was still present.

| Strain | Acetate (mM) |  |  | OD600 |  |  | Acetate/OD |  |  |
| --- | --- | --- | --- | --- | --- | --- | --- | --- | --- |
|  | time=3h | time=4h | time=5h | time=3h | time=4h | time=5h | time=3h | time=4h | time=5h |
| A1-C2 | 0.549 | 0.946 |  | 0.157 | 0.370 | 0.513 | 3.50 | 2.56 |  |
| A1-F5 | 0.528 | 0.912 |  | 0.152 | 0.323 | 0.509 | 3.47 | 2.82 |  |
| A1-D5 | 0.554 | 0.969 |  | 0.158 | 0.347 | 0.507 | 3.51 | 2.79 |  |
| A1-F6 | 0.537 | 0.934 |  | 0.154 | 0.320 | 0.507 | 3.49 | 2.92 |  |
| A1-E6 | 0.536 | 0.959 |  | 0.156 | 0.339 | 0.509 | 3.44 | 2.83 |  |
| A2-B6 | 0.736 | 1.237 |  | 0.223 | 0.528 | 0.530 | 3.30 | 2.34 |  |
| A2-D5 | 0.725 | 1.327 |  | 0.205 | 0.495 | 0.506 | 3.54 | 2.68 |  |
| A2-E9 | 0.648 | 1.258 |  | 0.196 | 0.464 | 0.490 | 3.31 | 2.71 |  |
| A2-E6 |  | 0.847 | 1.278 | 0.105 | 0.264 | 0.514 |  | 3.21 | 2.49 |
| A2-G7 | 0.745 | 1.316 |  | 0.210 | 0.503 | 0.522 | 3.55 | 2.62 |  |
| A3-E7 | 0.568 | 1.03 |  | 0.173 | 0.370 | 0.505 | 3.28 | 2.78 |  |
| A3-G11 | 0.594 | 1.099 |  | 0.177 | 0.401 | 0.495 | 3.36 | 2.74 |  |
| A3-D12 | 0.695 | 1.313 |  | 0.213 | 0.465 | 0.492 | 3.26 | 2.82 |  |
| A3-B12 | 0.86 | 1.247 |  | 0.249 | 0.530 | 0.523 | 3.45 | 2.35 |  |
| A3-G12 |  | 0.791 | 1.286 | 0.095 | 0.241 | 0.482 |  | 3.28 | 2.67 |
| MG1655 | 0.441 | 0.733 | 1.091 | 0.114 | 0.198 | 0.327 | 3.87 | 3.70 | 3.34 |
| IR1 | 0.572 | 1.068 |  | 0.176 | 0.378 | 0.505 | 3.25 | 2.83 |  |

**Supplementary Table 3:** Mutations identified in the genome of strain IR1.

| position | mutation | annotation | gene | description |
| --- | --- | --- | --- | --- |
| 2,760,454 | C→A | intergenic<br>(+60 / -93) | yfjI → / → yfjJ | CP4-57 prophage; uncharacterized protein/CP4-57 prophage; uncharacterized protein |
| 2,999,264 | G→A | Q210* (<u>C</u>A<br>A→<u>T</u>AA) | ygeR ← | LytM domain-containing M23 family putative peptidase; septation lipoprotein |
| 3,026,033 | Δ1 bp | coding (268<br>/ 1320 nt) | guaD → | guanine deaminase |
| 3,815,859 | Δ82 bp | pseudogene<br>(610-691 /<br>716 nt) | rph ← pyrE | ribonuclease PH (defective); enzyme; degradation of RNA; RNase PH |
| 3,984,420 | T→A | T530S (<u>A</u>C<br>C→<u>T</u>CC) | aslA ← | putative Ser-type periplasmic non-aryl sulfatase |

**Supplementary Table 4:** Identifying growth rates of E. coli which allow full respiration. The two rows at the bottom of the table show that for the two strains used in this study the growth rate reduction on acetate is much bigger compared to the growth rate reduction needed for cells to fully respire (top part of the table).

| Reference | Figure /table | Strain | Culture method | Medium | $\mu$ at which cells still show full respiration | $\mu_{\max}$ /highest D used in paper | Fraction of $\mu_{\max}$ at which cells still show full respiration |
| --- | --- | --- | --- | --- | --- | --- | --- |
| Nanchen (2006)<sup>6</sup> | Fig. 3 | MG1655 (l- F- rph-1Fnr+; DSMZ) | chemostat | M9 minimal | 0.4 | 0.7 | 0.57 |
| Holms (1996)<sup>7</sup> | Table 7 | ML308 = ATCC15224 | chemostat |  | 0.72 | 0.94 | 0.77 |
| Valgepea (2010)<sup>8</sup> | Fig. 2 | K12 MG1655 (l- F- rph-1Fnr+; DSMZ) | chemostat | defined minimal | 0.27 | 0.46 | 0.59 |
| Renilla (2012)<sup>9</sup> | Fig 4 | BW25113 a K12-derivative | chemostat | defined minimal | 0.5 | 0.7 | 0.71 |
| Meyer (1984)<sup>10</sup> | Fig. 1 | defined medium | chemostat |  | 0.35 | 0.54 | 0.65 |
| Basan (2015)<sup>11</sup> | Fig. 1 | K12=MG1655 | batch | minimal | 0.76 | 1.26 | 0.60 |
| | | | | | $\mu_{\max}$ on acetate | $\mu_{\max}$ on glucose | Fraction of $\mu_{\max}$ |
| This Study |  | MG1655 | batch | M9 minimal | 0.22 | 0.84 | 0.26 |
| This Study |  | IR1 - evolved | batch | M9 minimal | 0.29 | 1.1 | 0.26 |

**Supplementary Table 5:** Growth rates on acetate and glucose, and the ratio between them for a number of different organisms and strains. ^a^Strain not able to grow on glucose, therefore malate was taken as a reference high quality carbon source. ^b^Strain unable to grow on glucose, so no ratio was calculated.

| Organism | $\mu$ on acetate | $\mu$ on glc | $\mu_{ac}/\mu_{glc}$ | Reference |
| --- | --- | --- | --- | --- |
| *E. coli* MG1655 | 0.22 | 0.84 | 0.28 | This study |
| *E. coli* IR1 | 0.29 | 1.10 | 0.31 | This study |
| *Yarrowia lipolytica* | 0.28 | 0.41 | 0.68 | Robak (2007)<sup>12</sup> |
| *Corynebacterium glutamicum* | 0.28 | 0.32 | 0.88 | Wendisch et al. (2000)<sup>13</sup> |
| *Rhodobacter capsulatus* | 0.29 | 0.42<sup>a</sup> | 0.71 | Willison (1988)<sup>14</sup> |
| *E. coli* B/r strain NF790 | 0.30 | 0.89 | 0.34 | Andersen&Meyenburg (1980)<sup>15</sup> |
| *S. cerevisiae* AG1-7 | 0.12 | 0.31 | 0.39 | Johnston et al. (1979)<sup>16</sup> |
| *E. coli* BW25113 | 0.29 | 0.60 | 0.48 | Volkmer&Heinemann (2011)<sup>17</sup> |
| *Neurospora crassa* | 0.41 | 0.51 | 0.80 | Alberghina et al. (1975)<sup>18</sup> |
| *Acinetobacter schindleri*<sup>b</sup> | 0.91 |  |  | Sigala (2019)<sup>19</sup> |

